# Neutrophil remodeling is associated with human meibomian gland dysfunction and enables IFN-γ- and PAD4-dependent gland obstruction in mice

**DOI:** 10.64898/2026.08.19.744915

**Authors:** Cole J. Beatty, Symon Ma, Oleg Kolupaev, John B. Cart, Hazem M. Mousa, Rose Mathew, Daniel Floyd, John M. Fallon, Kevin R. Kipp, Justyna Resztak, Zhuoya Wan, Areej Ammar, Sejiro Littleton, Chen Yu, Eden M. Jacob, Elina Regan, Shruti Mistry, Agnes Acevedo Canabal, Audrey Nguyen, Joan Kalnitsky, Katherine S. Held, Victor L. Perez, Daniel R. Saban

## Abstract

Meibomian gland dysfunction (MGD), a disorder of the eyelid’s modified sebaceous glands, is the leading cause of dry eye disease and ocular surface morbidity, yet the immune mechanisms driving gland obstruction remain poorly defined. In a cross-sectional study of 66 patients with ocular surface inflammation, we used meibography and spectral flow cytometry of tear washes to identify a disease-associated, remodeled neutrophil state whose abundance is associated with gland atrophy. Using single-cell transcriptomics in a murine model of immune-mediated MGD, we revealed a disease-associated neutrophil state that exhibited ocular surface-enrichment, CD14 and ICAM-1 expression, and elevated IFN-γ response and inflammatory signatures. Spatial transcriptomics localized IFN-γ signaling and neutrophil migration signatures to the periglandular compartment. The remodeled neutrophils exhibited PAD4-dependent histone citrullination, with *Padi4* deletion reducing NET-associated obstructive plugging, thus identifying PAD4-dependent NETotic activity as their disease-producing output. Inhibition of IFN-γ signaling phenocopied *Padi4* deficiency, yet combined disruption of these pathways provided no additive protection, indicating that IFN-γ and PAD4 function as separable required inputs. Remodeled neutrophils accumulated under both conditions, uncoupling disease severity from cell abundance alone. Our findings support immune-mediated obstructive MGD as a mechanistic endotype driven by the IFN-γ- and PAD4-dependent effector output of a remodeled neutrophil state.

**GRAPHICAL ABSTRACT:** 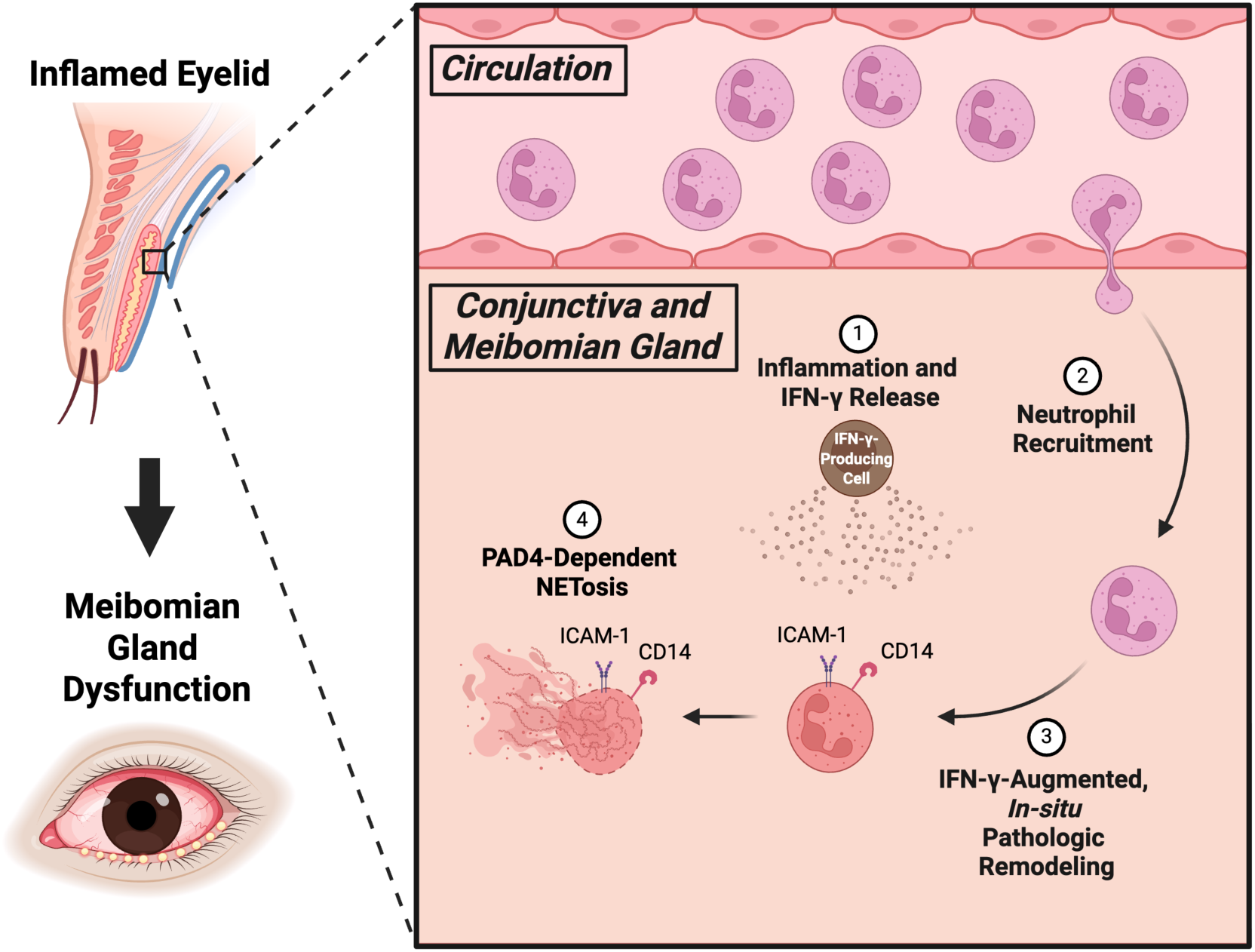

## INTRODUCTION

Meibomian glands are modified sebaceous glands lining the eyelid that secrete lipids essential for lubricating the ocular surface. Dysfunction of these glands, termed meibomian gland dysfunction (MGD), is the leading cause of dry eye disease and one of the most common conditions encountered in ophthalmology clinics worldwide (1, 2). The most common form is obstructive MGD, in which gland orifices become blocked, leading to ductal dilation, acinar atrophy, and permanent gland dropout (3–6). Once atrophy is established, gland loss is irreversible, and the progressive ocular surface deterioration which follows can threaten vision (7–10). Despite the global burden of this disease, there is no FDA-approved disease-modifying pharmacotherapy for obstructive MGD, and treatments largely remain palliative (1, 11).

The classical view of MGD pathogenesis emphasizes epithelial, ductal, lipid, hormonal, and age-associated mechanisms, with inflammation long being considered a secondary feature of MGD (5). However, this framework does not fully explain why MGD is a prominent and often severe complication of ocular surface inflammatory disorders (OSIDs), including ocular graft-versus-host disease (oGVHD), Sjögren’s disease, Stevens-Johnson syndrome, and chronic allergic eye disease, including atopic and vernal keratoconjunctivitis and blepharokeratoconjunctivitis (12–16). In the allergic eye disease (AED) model, an immune-mediated model of ocular surface inflammation that develops obstructive MGD, Reyes et al. showed that IL-17A-producing CD4^+^ T cells promote the recruitment of neutrophils that obstruct meibomian gland orifices, and that tear neutrophil abundance tracks with MGD severity across OSID diagnoses (17). Consistent with this finding, Perez et al. showed that oGVHD associated with MGD is linked to neutrophil increases in human tears and elevation of IL-17A-producing CD4^+^ T cells in the draining lymph nodes of mice (13). Similarly, Postnikoff et al. identified increases in granulocyte to lymphocyte ratios in the ocular surface washes of patients with dry eye disease (18). Neutrophil infiltration and IL-17/IL-23 inflammatory pathway activation also arise in the Pinkie/RXRα mutant model of inflammatory dry eye disease (19, 20) that develops meibomian gland dysfunction (21), as well as in a non-immune model of MGD with *Awat2* deficiency (22, 23). These observations suggest that specific types of inflammation, including neutrophil responses, may represent a convergent disease-modifying pathway across distinct etiologies of MGD (24). Relatedly, neutrophils are increasingly recognized as heterogeneous cells whose distinct transcriptional states can drive disease-specific functional outputs in inflammatory and neoplastic settings (25–31). However, whether neutrophil remodeling occurs in MGD, and whether specific neutrophil states shape glandular obstruction, remains unknown.

Despite the co-occurrence of inflammation and obstructive MGD across OSIDs, the immune mechanisms that link ocular surface inflammation to glandular obstruction remain poorly defined. We asked whether disease-associated neutrophil states mediate this transition and investigated what upstream signals govern their pathogenic activity. In a cross-sectional study of 66 patients with ocular surface inflammation, we identified a remodeled tear neutrophil state whose abundance is associated with MGD severity. Similarly, in a murine model of immune-mediated obstructive MGD, we resolved a corresponding neutrophil state, hereby labelled PMN4, and identified IFN-γ and PAD4 as separable inputs required for PMN4-induced obstructive MGD. Together, these findings support immune-mediated obstructive MGD as a mechanistic endotype driven by the IFN-γ- and PAD4-dependent effector output of a remodeled neutrophil state.

## RESULTS

### Patient MGD Severity is Associated with Tear Neutrophil Remodeling

Tear neutrophil abundance was previously shown to correlate with deterioration of meibum quality in patients with MGD (17), but the broader immune landscape of patient tears and the phenotypic heterogeneity of tear neutrophils had not been characterized. To address this, we enrolled 66 patients with ocular surface inflammation, collected tear washes, and performed infrared meibography (32) to score meibomian gland atrophy as a measure of MGD severity (Table S1, Fig. 1a). Tear cells were immunophenotyped using a 36-parameter spectral flow cytometry panel first published by Yu et al. for broad leukocyte characterization of human samples (13, 33) and analyzed alongside healthy donor blood as a phenotypic reference (Fig. 1a, Table S2). Major leukocyte populations were identified in both sample types (Fig. 1b,c; Fig. S1a, Table S3), with neutrophils disproportionately enriched in tears relative to leukocyte populations in reference blood samples (Fig. 1d). Within the tear neutrophil compartment, we observed marked phenotypic heterogeneity. A subset of tear neutrophils exhibited elevated forward scatter (FSC), side scatter (SSC), CD16, CD95, CXCR3, CD45, and CD11b, together with reduced CD127 (Fig. 1e). Because increased CD11b and FSC have been reported following neutrophil activation by inflammatory stimuli (34–36), this phenotype is consistent with an inflammatory-like remodeled tear neutrophil state.

**Figure 1.**
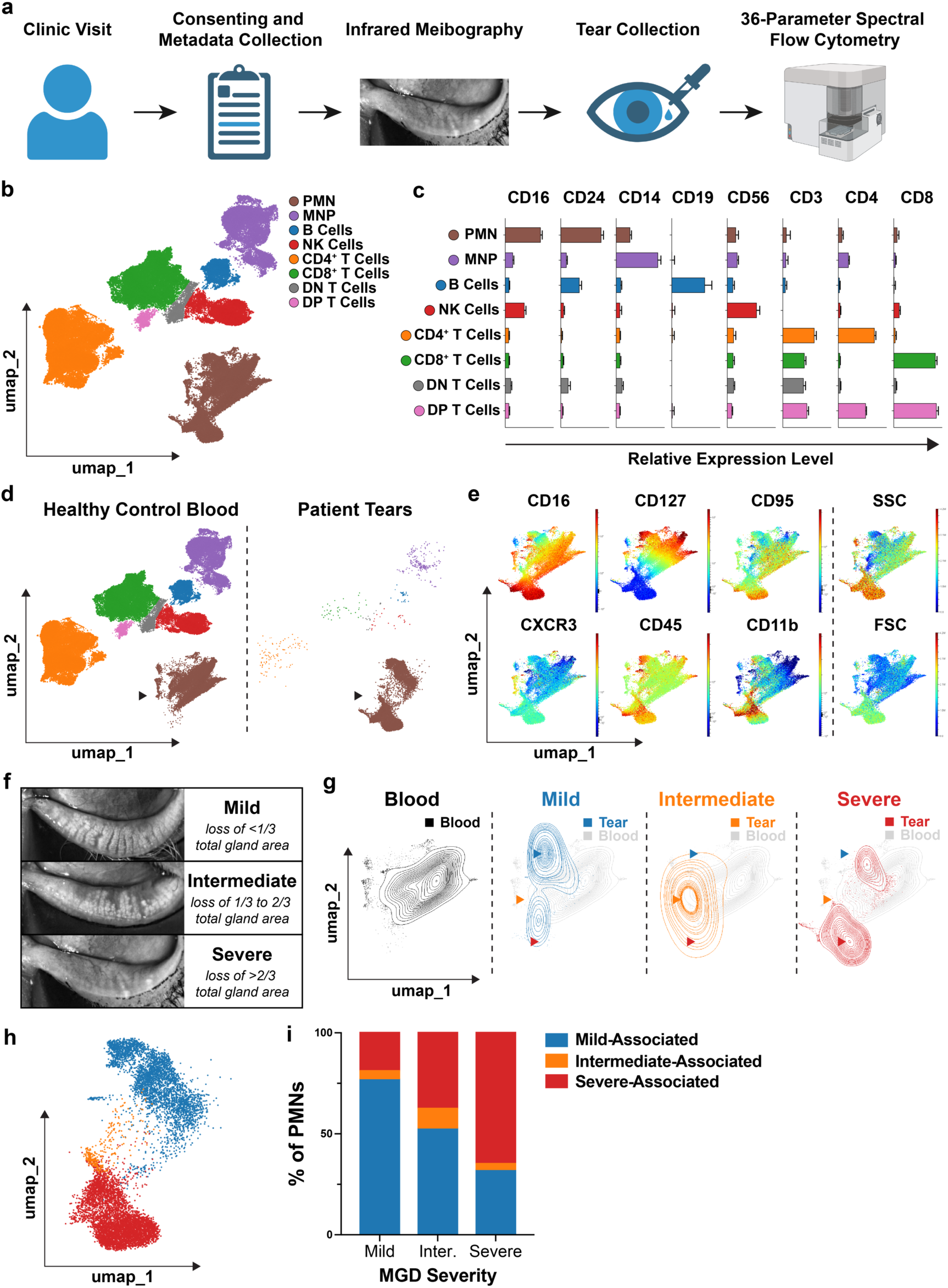
A remodeled, inflammatory-like tear neutrophil phenotype is enriched in patients with severe MGD. **(A)** Experimental design overview. N = 66 patients. **(B)** UMAP of concatenated normal control blood and tear leukocytes stained with 36-parameter flow cytometry panel. **(C)** Established protein marker expression patterns for each cell population, supporting UMAP annotations. Bars represent the mean fluorescence intensity for each respective marker as detected from individual patients. **(D)** Leukocyte UMAP separated based on origination from healthy control blood or patient tear sample. Black arrows emphasize neutrophils. **(E)** Visualizing concatenated tear and blood neutrophil marker expression, demonstrating heterogeneity based on established inflammatory markers. **(F)** Representative infrared meibography images for mild, intermediate, and severe MGD severity scoring determinations. **(G)** Comparing neutrophil heterogeneity by overlaying UMAPs of tear neutrophils from patients with mild, intermediate, and severe MGD over a healthy blood control neutrophil UMAP. **(H)** FlowSOM clustering of tear leukocytes flow data for unbiased heterogeneity analysis. UMAP shows the 3 largest neutrophil clusters. **(I)** Group summary plots showing proportion of neutrophils in each of the 3 largest neutrophil FlowSOM clusters by patient MGD severity. Plots show mean + SEM.

We next asked whether the severity of meibomian gland atrophy was associated with this tear neutrophil phenotype. Patients with severe MGD showed a shift in tear neutrophil phenotype toward higher FSC, SSC, CD16, CD95, CXCR3, CD45, CD11b, and reduced CD127 relative to patients with mild or intermediate MGD (Fig. 1e,f,g). In agreement with the observed phenotypic shift, unsupervised FlowSOM analysis identified three major PMN clusters whose distribution shifted with MGD severity (Fig. 1h,i), consistent with the development of distinct tear neutrophil phenotypes across the MGD severity spectrum. Although the trend was consistent, patient-to-patient variability was observed, with some patients exhibiting a lack of tear PMNs regardless of MGD status (Table S4) as previously reported by Reyes et al. (17). Together, these data indicate that meibomian gland atrophy severity in patients with ocular surface inflammation is associated with increased abundance of a remodeled, inflammatory-like tear neutrophil state.

### A Disease-Associated, Ocular Surface-Enriched Neutrophil Remodeled State in Murine Obstructive MGD

To characterize neutrophil remodeling in a mouse paradigm of immune-mediated obstructive MGD, we performed single-cell RNA sequencing (scRNA-seq) on blood, conjunctiva, and tears from AED and naïve mice. The AED model captures cardinal features of immune-mediated obstructive MGD, including neutrophil recruitment and meibomian gland orifice obstruction (14, 17) (Fig. 2a). Leukocytes were isolated via fluorescence activated cell sorting (FACS) before RNA capture; conjunctiva samples were prepared as a 1:1 mixture of Ly6G^+^ neutrophils and Ly6G^-^ leukocytes to ensure representation of both neutrophil and non-neutrophil immune compartments (Fig. S2a). Unsupervised clustering resolved 14 clusters, which were annotated using canonical markers (Fig. 2b,c; Table S5). Focusing on neutrophils, we identified four transcriptionally distinct populations, PMN1, PMN2, PMN3, and PMN4 (Fig. 2b,c). PMN4 was almost exclusively present in AED mice and absent from naïve controls (Fig. 2d), establishing it as a disease-associated neutrophil state. In contrast to PMN1-3, which were well represented in blood, PMN4 was enriched in conjunctiva and tears and largely absent from the systemic circulation (Fig. 2d), suggesting that it is shaped within the ocular inflammatory microenvironment.

**Figure 2.**
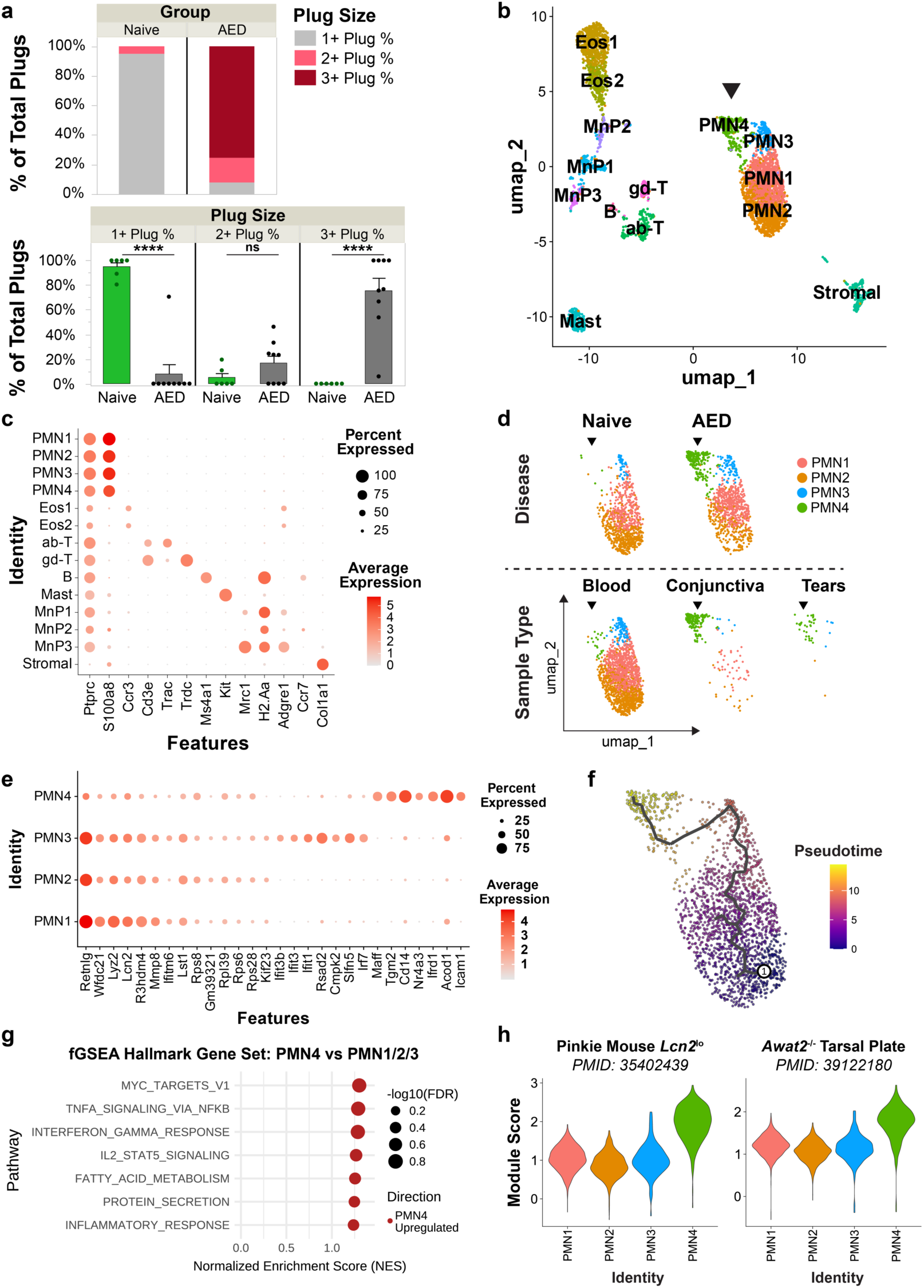
PMN4 Is a disease-associated, ocular surface-enriched neutrophil remodeled state in murine obstructive MGD. **(A)** MG plug scoring for WT naive and AED mice. Stacked bar plot represents group summary plug scoring and individual bar plots display individual mouse values for each plug size. Dots represent respective plug size percentage for each mouse individually. N = 9 AED mice, N = 6 naive mice. Dots represent values for individual mice. **(B)** UMAP of scRNA-seq data from WT naïve and AED mouse blood, conjunctiva, and tears. Black arrow indicates PMN4 population of interest. PMN, polymorphonuclear neutrophils; Eos, eosinophils; MnP, mononuclear phagocytes; gd-T, γδ T cells; ab-T, αβ T cells **(C)** Dotplot showing expression of established marker genes for cellular annotation. **(D)** PMN1-4 split by group or sample type of origin. Black arrows indicate PMN4 population of interest. **(E)** Top 7 DEGs distinguishing each PMN cluster from other PMNs. **(F)** Monocle3 pseudotime analysis of PMNs beginning with initialization point at PMN2. **(G)** Hallmark Gene Set fGSEA pathway analysis evaluating differentially expressed gene pathways upregulated in PMN4 relative to PMN1-3. **(H)** Module scoring of PMN1-4 using DEGs which identify neutrophil subpopulations observed in published murine models of MGD, namely the Pinkie and *Awat2*^-/-^ mouse models. ****: P <0.0001 ; ns: not significant. ANOVA with Tukey’s HSD post-hoc test (A). Plots show mean + SEM.

Because PMN4 was uniquely disease-associated and ocular surface-enriched, we next asked what transcriptional programs distinguished it from the other neutrophil populations and whether it represented a terminal remodeled state. PMN4 was distinguished from PMN1-3 by upregulation of inflammatory and activation-associated genes, including *Il1a*, *Rab20*, *Ccl3*, *Ccl4*, *Tnf*, *Icam1*, and *Cd14* (Fig. 2e, S2b; Table S6, S7). Pseudotime trajectory analysis (37, 38), initialized at PMN2 based on its enrichment in naïve samples and blood, as well as expression of neutrophil immaturity markers including *Mmp8*, *Mmp9*, and *Cxcr2* (26, 27, 30) (Fig. S2c,d), identified PMN4 as the terminal population (Fig. 2f), consistent with a disease-associated remodeling trajectory. Fast gene set enrichment analysis (fGSEA) (39) comparing PMN4 to PMN1-3 using the Mouse Hallmark Gene Set (40, 41) revealed enrichment of Interferon Gamma Response, TNFα Signaling via NFκB, and Inflammatory Response programs, alongside fatty acid metabolism and protein secretion pathways (Fig. 2g). To evaluate whether AED-defined neutrophil states are represented across MGD etiologies, we applied gene modules (42) from DEGs that define neutrophils enriched in *Awat2*-deficient mice (22, 23) and *Lcn2*^lo^ neutrophils from Pinkie/RXRα mutant mice (19, 20) to PMN1-4 (Table S8). This analysis identified PMN4-like neutrophils in both datasets (Fig. 2h), suggesting that the PMN4 remodeled state may not be restricted to immune-initiated disease. Together, these data identify PMN4 as a disease-associated, ocular surface-enriched neutrophil state in MGD and support the use of the AED model as a murine system to test how neutrophil remodeling contributes to meibomian gland obstruction.

### PMN4 Is the Principal PAD4-Dependent Histone-Citrullinated Neutrophil State in Murine Obstructive MGD

We next asked whether PMN4 plays a causal role in obstructive MGD. Taking a clue from Mahajan et al., who demonstrated that PAD4-dependent neutrophil extracellular trap formation (NETosis) drives meibomian gland orifice obstruction in the AED model through formation of aggregated NETs (43), we asked whether PMN4 is the principal neutrophil population linked to PAD4-dependent histone citrullination and gland obstruction. We compared WT AED mice to *Padi4*-deficient (44, 45) AED mice and evaluated MGD severity and PMN4 abundance. *Padi4* deficiency reduced obstructive plugging compared with WT AED controls (Fig. 3a), extending prior evidence that PAD4-dependent NETosis (46, 47) contributes to gland obstruction in this model (43). To determine whether this protection reflected changes in neutrophil composition, we performed flow cytometry on blood and conjunctiva using a PMN4-focused spectral flow cytometry panel incorporating ICAM-1, CD14, CXCR2, and CD62L (Table S9). Total conjunctival neutrophil counts were not significantly different between groups, but the proportion of conjunctival neutrophils with a PMN4 phenotype was significantly elevated in *Padi4*-deficient AED mice compared with WT AED controls (Fig. 3b-d, Fig. S3a). Thus, *Padi4* deficiency separated PMN4 remodeling from gland obstruction: PMN4s accumulated, but obstructive plugging was reduced. These findings indicate that PAD4 is necessary for gland obstruction and raise the question of whether PMN4 is the predominant neutrophil population executing PAD4-dependent citrullination at the meibomian gland.

**Figure 3.**
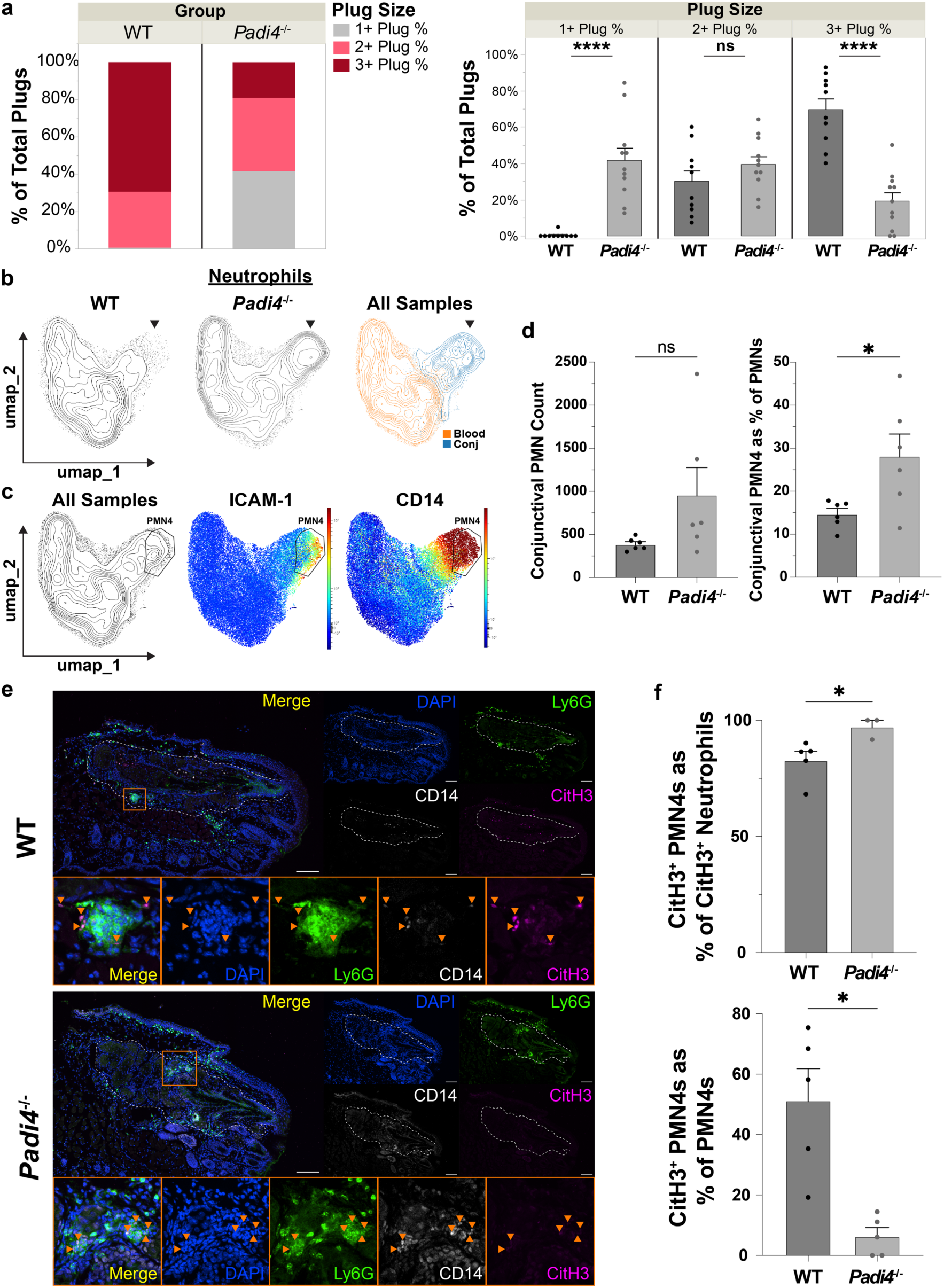
PMN4 is the predominant neutrophil population executing PAD4-dependent histone citrullination at the meibomian gland. **(A)** MG plug scoring for WT AED and *Padi4*^-/-^ AED mice. Stacked bar plot represents group summary plug scoring and individual bar plots display individual mouse values for each plug size. Dots represent respective plug size percentage for each mouse individually. N = 10 WT AED, N = 11 *Padi4*^-/-^ AED mice. **(B)** Concatenated flow cytometry UMAPs for neutrophils isolated from blood and conjunctiva of WT AED and *Padi4*^-/-^ AED mice. Black arrows indicate neutrophil population enriched in *Padi4*^-/-^ AED conjunctiva. **(C)** Concatenated neutrophil flow cytometry UMAPs with visualization of *Padi4*^-/-^-enriched PMN4 population gating and key PMN4 DEG-guided markers ICAM-1 and CD14. **(D)** Quantifications of flow cytometry data from WT AED and *Padi4*^-/-^ AED conjunctiva, specifically PMN count and the percentage of PMNs in the *Padi4*^-/-^ -enriched PMN4 gate. Dots represent values for individual mice. **(E)** Representative staining of Ly6G (green), CD14 (white), CitH3 (pink) in the eyelids of WT AED and *Padi4*^-/-^ AED mice. Orange arrows indicate CD14^+^ Ly6G^+^ PMN4s. Orange boxes indicate areas in and around the meibomian glands that have been magnified below the full images for closer evaluation. Scale bars: 100 μm. **(F)** Quantification of the percentage of CitH3^+^ Ly6G^+^ neutrophils coexpressing CD14 and the percentage of CD14^+^ Ly6G^+^ PMN4s coexpressing CitH3 in WT AED and *Padi4*^-/-^ AED stained eyelids. Dots represent values for individual mice. Data were collected from 2-3 independent experiments. *: P < 0.05 ; ****: P <0.0001 ; ns: not significant. Repeated-measures ANOVA with Tukey’s HSD post-hoc test (A), Welsch’s T tests (D), Mann-Whitney test and Welsch’s T test, respectively (F). Plots show mean + SEM.

To evaluate whether PMN4 exhibits PAD4-dependent histone citrullination at the meibomian gland, we performed immunofluorescence imaging of eyelids from WT and *Padi4*-deficient AED mice using Ly6G, CD14, and citrullinated histone H3 (CitH3) (47) as markers of neutrophils, PMN4 identity, and PAD4-dependent chromatin citrullination, respectively (Fig. 3e). In WT AED eyelids, CD14^+^ Ly6G^+^ PMN4s were the predominant CitH3^+^ neutrophil population in the periglandular space, with greater than 75% of CitH3^+^ neutrophils co-expressing CD14 (Fig. 3e,f). CitH3 staining was evident in approximately 50% of PMN4s in WT AED lids, consistent with citrullination reflecting an activation state within the PMN4 population (Fig. 3e,f). Moreover, CitH3 staining in the PMN4s was markedly reduced in *Padi4*-deficient AED eyelids, confirming that the citrullination signal was PAD4-dependent (Fig. 3e,f). Together with prior evidence that PAD4-dependent aggregated NETs obstruct meibomian gland orifices in AED-associated MGD, these data identify PMN4 as the principal neutrophil state exhibiting PAD4-dependent histone citrullination at the meibomian gland and support PAD4-dependent NETotic effector activity as the obstruction-producing output of the PMN4 remodeling state.

### IFN-γ Signaling and Neutrophil Migration Programs Are Enriched in the Periglandular Compartment in Murine Obstructive MGD

To identify tissue-level signals that may shape PMN4 remodeling and its obstruction-producing activity at the meibomian gland, we performed whole-transcriptome spatial RNA sequencing using GeoMx digital spatial profiling on eyelids from AED and naïve mice (Fig. 4a,b). Regions of interest (ROIs) were selected in collaboration with histopathologists to capture three anatomically distinct compartments: the meibomian gland duct, defined as pan-cytokeratin (panCK)-positive regions with ductal morphology; acini, defined as panCK-positive acinar regions with acinar morphology; and the inter-acini compartment, defined as the panCK-negative region surrounding the acini (Fig. 4a,b). Following quality control processing (Fig. S4a-c), unsupervised analysis separated AED and naïve ROIs, with the most pronounced disease-associated transcriptional differences observed in the acinar and inter-acini compartments (Fig. 4c). Relative to naive controls, AED ROIs showed broad transcriptional changes including upregulation of *Lcn2*, a marker previously associated with neutrophil infiltration in MGD (22), and *Cxcl10*, an IFN-γ-induced chemokine (48) (Fig. 4d, Fig. S4d, Table S10-12).

**Figure 4.**
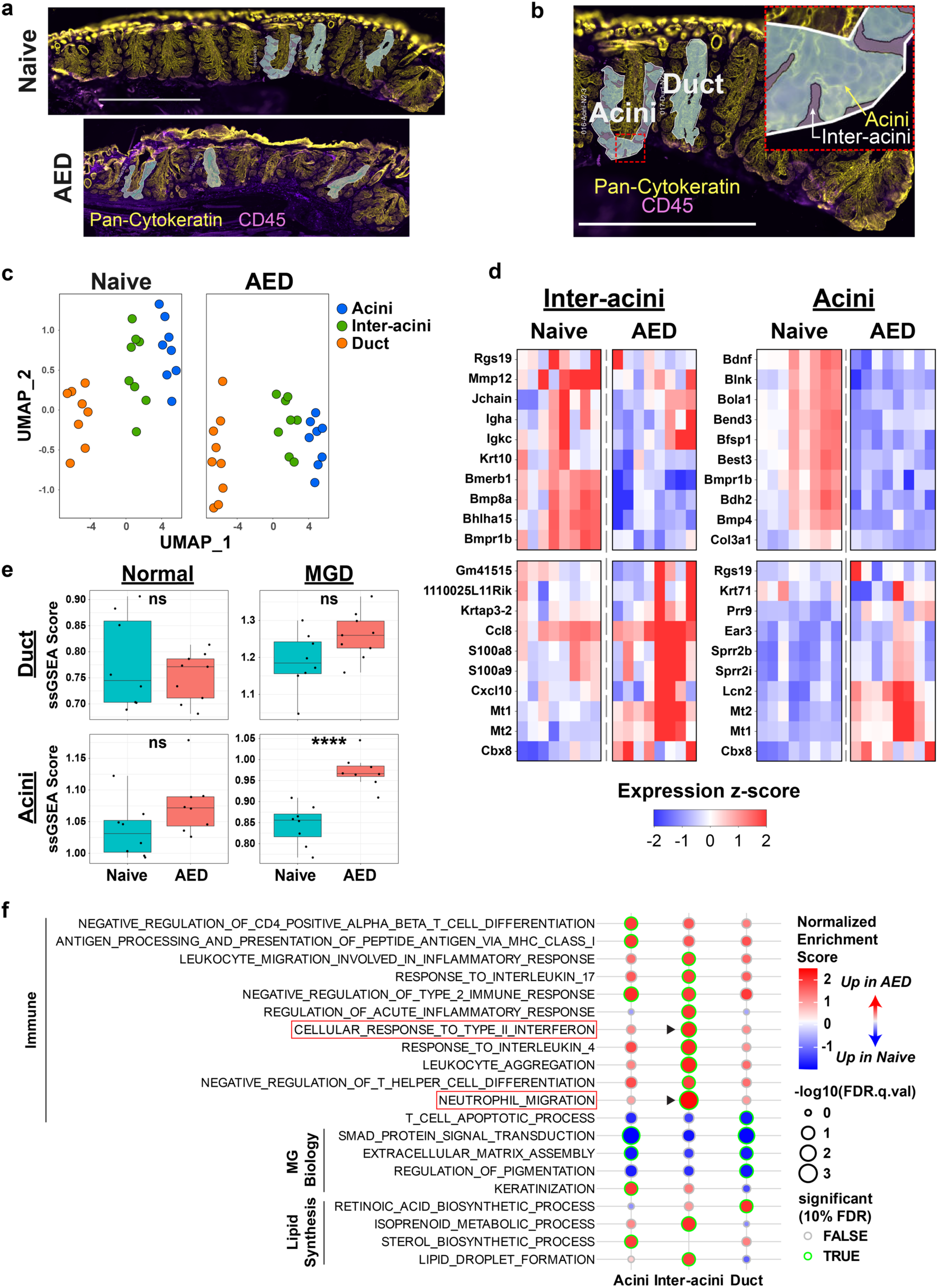
Inflammatory pathways that drive MGD, including neutrophil recruitment, exist in close spatial association with IFN-γ responses. **(A)** Representative images from WT naive and AED eyelid sections used for GeoMx analysis. N = 2 WT AED and N = 2 WT naive mice. Scale bar: 1 mm. **(B)** ROI generation schematic. N = 8-9 duct ROIs and 8 acini/inter-acini ROIs per group. Scale bar: 1 mm. **(C)** GeoMx spatial transcriptomics UMAP for ROIs selected from the acini, inter-acini, and duct of WT naive and AED mice. Dots represent values for individual ROIs. **(D)** Heatmaps displaying DEGs that distinguish naive and AED eyelids by ROI for inter-acini and acini compartments. **(E)** ssGSEA of WT naive and AED acini and duct ROI transcriptomic signatures using published scRNA-seq DEG signatures that distinguish healthy B6 and *Awat2*^-/-^ MGD acini and ducts. Dots represent values for individual ROIs. **(F)** GSEA comparing WT AED and naive eyelids by ROI compartment. Red boxes highlight key inflammatory pathways related to neutrophils and IFN-γ. ****: P<0.0001; ns: not significant. T-test (E).

To ask whether immune-mediated gland obstruction produces transcriptional changes relevant to other MGD etiologies, we performed single-sample gene set enrichment analysis (ssGSEA) (46) using published DEG signatures which distinguish healthy and *Awat2*-deficient meibocytes and ducts (22) (Table S13). AED acini, but not duct, showed significant enrichment for the *Awat2*-deficient MGD meibocyte signature compared with naïve controls, while enrichment against the healthy meibocyte signature was not significant in either compartment (Fig. 4e), suggesting that this transcriptional convergence with non-immune MGD is specific to the acinar compartment and to the disease-associated gene signature.

We next asked which pathways were enriched in AED eyelids and whether they were spatially associated with neutrophil recruitment programs. GSEA using the Gene Ontology Biological Process gene sets (49, 50) on the DEGs which distinguish AED and naïve ROIs by compartment showed that immune pathway enrichment was largely restricted to the inter-acini compartment, while meibomian gland biology and lipid synthesis pathways were enriched in the acinar and ductal compartments (Fig. 4f), indicating spatially compartmentalized disease programs. Within the inter-acini compartment, neutrophil migration and IL-17 response pathways were enriched in the inter-acini region of AED mice (Fig. 4f), consistent with prior evidence that IL-17A-driven neutrophil recruitment leads to gland obstruction in this model (17). Cellular response to IFN-γ was also enriched in the inter-acini compartment, in close spatial association with the neutrophil migration signature (Fig. 4f). Although elevated IFN-γ responses were previously noted in the draining lymph nodes of AED mice (17), their spatial enrichment within the periglandular compartment was not established previously. Together, these data indicate that IFN-γ signaling and neutrophil migration programs are enriched in the inter-acini space surrounding the meibomian gland in AED-associated MGD, raising the question of whether IFN-γ contributes to MGD pathogenesis through its action on the PMN4-associated obstructive program.

### IFN-γ and PAD4 are Separable Required Inputs to PMN4-Mediated Obstruction in Murine Obstructive MGD

To test whether IFN-γ contributes to obstructive MGD, we inhibited IFN-γ signaling by two independent strategies: global *Ifngr1* deletion (51) and systemic antibody-mediated IFN-γ blockade (52, 53) in WT AED mice during the challenge phase. Both genetic and pharmacologic loss of IFN-γ signaling reduced obstructive plugging compared with controls (Fig. 5a,b), establishing that IFN-γ is required for MGD immunopathogenesis in the AED model. To ask whether IFN-γ and PAD4-dependent NETotic activity act through independent or shared mechanisms, we performed systemic IFN-γ blockade in both WT and *Padi4*-deficient AED mice and compared MGD severity across groups (Fig. 5c). IFN-γ blockade in *Padi4*-deficient mice did not reduce obstructive plugging beyond IFN-γ blockade alone (Fig. 5d, S5a), indicating that IFN-γ and PAD4 operate within a shared obstruction-producing mechanism. To determine whether IFN-γ blockade influenced PMN4 abundance, we performed flow cytometry on conjunctiva and tarsal plate from each group. PMN4 counts were elevated following IFN-γ blockade, mirroring the accumulation observed in *Padi4*-deficient AED mice, and PMN4 counts were further increased when IFN-γ blockade and *Padi4* deficiency were combined (Fig. 5e), suggesting IFN-γ may not be necessary for PAD4 activity despite both being required for obstruction. To determine whether IFN-γ was required for PAD4-dependent histone citrullination in PMN4s, we performed immunofluorescence imaging of eyelids from each group using Ly6G, CD14, and CitH3 (Fig. 5f). PMN4s in the IFN-γ blockade condition retained CitH3 staining (Fig. 5f), demonstrating that PAD4-dependent histone citrullination can proceed despite IFN-γ blockade. Together, these data indicate that IFN-γ is required for the obstruction-producing output of PMN4 but is dispensable for PMN4 accumulation and PAD4-dependent histone citrullination.

**Figure 5.**
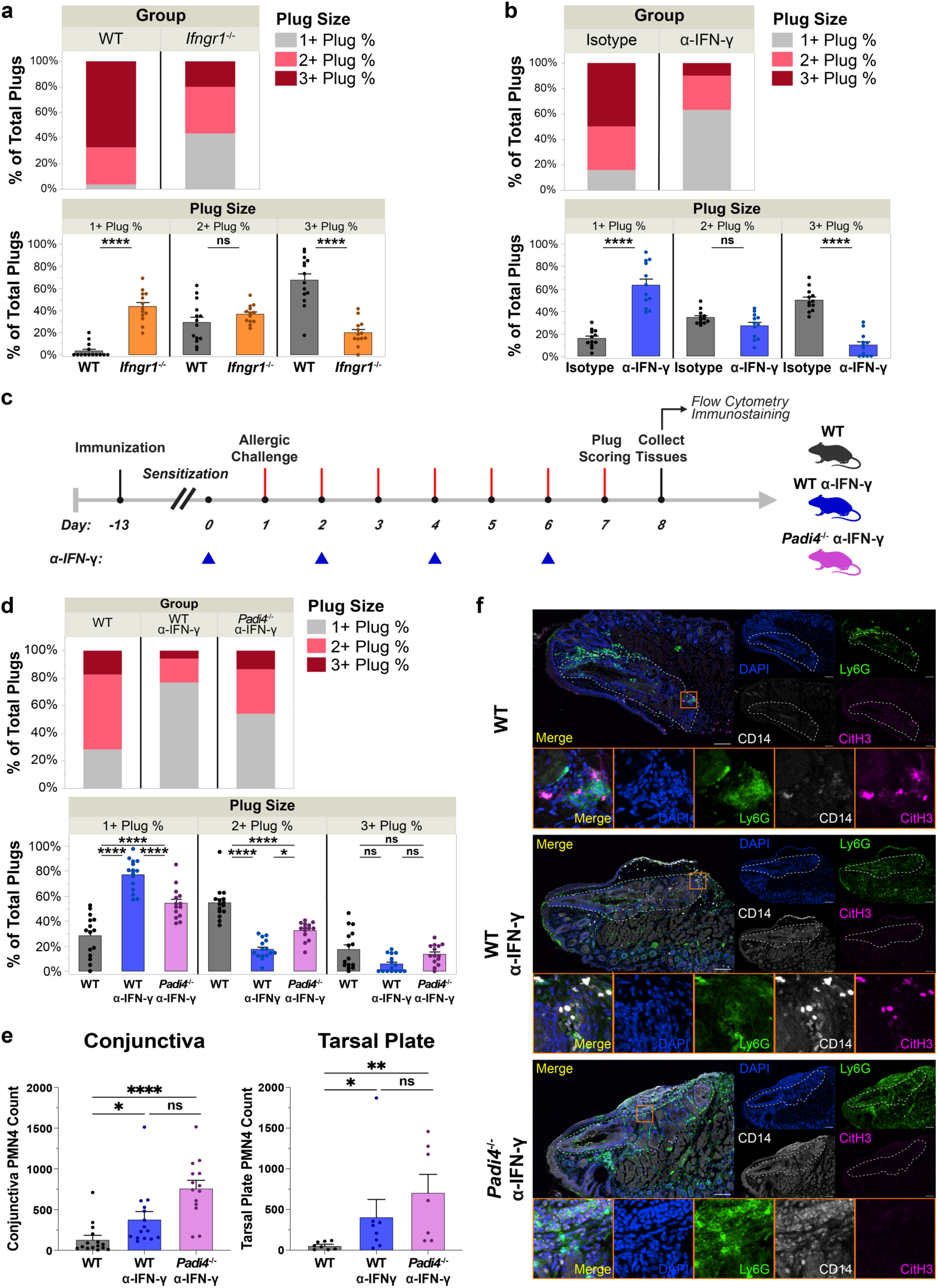
IFN-γ and PAD4 are separable inputs required for PMN4-mediated obstruction. **(A)** MG plug scoring for WT AED (N = 14) and *Ifngr1*^-/-^ AED (N = 13) mice. Stacked bar plot represents group summary plug scoring and individual bar plots display individual mouse values for each plug size. Dots represent respective plug size percentage for each mouse individually. **(B)** MG plug scoring for WT AED mice receiving systemic isotype control antibody (N = 12 mice) or systemic IFN-γ blockade antibody (N = 12 mice) injections. Stacked bar plot represents group summary plug scoring and individual bar plots display individual mouse values for each plug size. Dots represent respective plug size percentage for each mouse individually. **(C)** Experiment design overview. N = 15 WT AED, N = 15 WT AED receiving IFN-γ blockade, and N = 14 *Padi4*^-/-^ AED receiving IFN-γ blockade. **(D)** MG plug scoring for WT AED, WT AED with systemic IFN-γ blockade antibody instillation, and *Padi4*^-/-^ AED with systemic IFN-γ blockade antibody instillation. Stacked bar plot represents group summary plug scoring and individual bar plots display individual mouse values for each plug size. Dots represent respective plug size percentage for each mouse individually. **(E)** Flow cytometry counts of ICAM-1^+^ CD14^+^ PMN4s isolated from the conjunctiva and tarsal plate for each group. Dots represent values for individual mice. **(F)** Representative staining of Ly6G (green), CD14 (white), CitH3 (pink) in the eyelids of each group. Orange boxes indicate areas in and around the meibomian glands that have been magnified below the full images for closer evaluation. Scale bars: 100 μm. Data were collected from 2-3 independent experiments. *: P < 0.05 ; **: P < 0.01 ; ****: P <0.0001 ; ns: not significant. Repeated-measures ANOVA with Tukey’s HSD post-hoc test (A,B, D), Kruskal-Wallis test (E). Plots show mean + SEM.

We next evaluated whether IFN-γ can directly affect remodeling of neutrophils towards a PMN4 phenotype. Critically, PMN4 uniquely expressed both subunits of the functional IFN-γ receptor, *Ifngr1* and *Ifngr2* (Fig. S5b), indicating that PMN4 cells may be directly IFN-γ-responsive (54). However, evaluating the ability of IFN-γ to directly influence neutrophil remodeling in vivo was complicated by the complex inflammatory milieu present in the AED model (17) and numerous non-neutrophil IFN-γ-responsive cell populations present at the ocular surface (55, 56).

Therefore, we leveraged a reductionist approach using ex vivo stimulation of thioglycollate-elicited peritoneal neutrophils (57, 58). We observed that recombinant IFN-γ contributed to acquisition of a PMN4 surface phenotype (CD14^+^ ICAM-1^+^ CD62L^lo^ CXCR2^lo^, Fig. S5c) only alongside a secondary inflammatory signal, bacterial lipopolysaccharide (LPS) (59, 60), with IFN-γ alone being insufficient to influence PMN4 formation and LPS stimulation alone producing only partial PMN4 phenotype induction (Fig. S5d,e,f). Taken together, these ex vivo tests demonstrate that IFN-γ can directly influence PMN4 biology, thus further supporting a role for IFN-γ in PMN4-mediated AED immunopathogenesis. Cumulatively, the in vivo and ex vivo data together establish IFN-γ and PAD4 as separable inputs required for the obstruction-producing activity of PMN4, with absence of either input resulting in reduced gland obstruction despite recruited neutrophils still being remodeled to the PMN4 state.

## DISCUSSION

In the current study, we show that inflammation in MGD is not merely a nonspecific feature of gland dysfunction but a state-dependent mechanism capable of producing obstruction. Rather than identifying neutrophils simply as recruited effector cells, our data support a conceptual framework in which the ocular surface inflammatory environment remodels neutrophils into a disease-associated state that is competent, but not by itself sufficient, to obstruct the meibomian gland. This distinction is central to immune-mediated obstructive MGD. The abundance of remodeled neutrophils can be associated with disease severity, but obstruction requires two additional inputs: PAD4-dependent histone citrullination and a separable IFN-γ-dependent requirement. Blocking either input reduces obstruction. Three implications emerge from this model. First, tear neutrophil remodeling may provide a clinically informative marker of inflammatory MGD severity; second, the PMN4 state links MGD to broader paradigms of disease-associated neutrophil specialization; and third, IFN-γ and PAD4 define separable inputs that gate the obstruction-producing activity of this remodeled state.

In the human cohort, tear neutrophil remodeling provided information beyond the presence of ocular surface inflammation alone. Although the human data remain associative, an inflammatory-like (34–36) remodeled tear neutrophil phenotype marked by elevated FSC, SSC, CD16, CD95, CXCR3, CD45, CD11b, and reduced CD127 was disproportionately abundant in severe MGD, extending our prior work linking tear neutrophil abundance with meibum quality loss (17) and linking the human tear neutrophil state to the inflammatory program observed in PMN4s. Moreover, the presence of CXCR3, a receptor for IFN-γ-induced chemokines (61, 62), links the human tear neutrophil state to the IFN-γ-associated program observed in murine PMN4 and in the periglandular inflammatory compartment. The broader relevance of neutrophils to ocular surface disease is increasingly recognized, with NETs implicated in dry eye pathology and ocular surface disease (63–65). Interestingly, not all patients with severe MGD exhibited this remodeled phenotype, suggesting that tear neutrophil remodeling may identify an inflammatory subgroup of MGD not captured by gland atrophy severity alone. This heterogeneity creates an opportunity for biomarker development and patient stratification in inflammatory MGD.

More broadly, the PMN4 state places MGD within an emerging paradigm of disease-associated neutrophil specialization. Neutrophils are increasingly recognized as transcriptionally and functionally heterogeneous, with disease-specific states described in cancer, autoimmunity, and infection (27, 28, 66). Specifically, PMN4 shares features with these non-ocular populations, including an enrichment in the inflammatory tissue site, IFN-γ-responsive and inflammatory transcriptional programs, and acquisition of surface phenotypes linked to altered effector potential. For example, ICAM-1^+^ neutrophils from both patients and mice have been associated with inflammatory settings and have been reported to exhibit elevated effector functions including phagocytosis and NETosis (67–69). This parallels our finding that ICAM-1^+^ PMN4s are an inflammatory population that exhibits elevated PAD4 activity known to be related to MGD-driving NETosis (43). Moreover, in tumor, autoimmune, and infectious settings, type II interferon signaling can shape neutrophil states in a context-dependent manner, often requiring additional inflammatory cues to specify downstream function (66, 70, 71). This framework parallels our ex vivo finding that IFN-γ alone was insufficient to induce a PMN4 phenotype, whereas IFN-γ cooperated with LPS to enhance acquisition of this remodeled surface state. PMN4 may therefore represent an ocular surface example of a broader principle in that inflammatory tissues do not merely recruit neutrophils, but remodel them into specialized disease-associated states whose pathogenic outputs are determined by local signals.

Mechanistically, our findings establish IFN-γ and PAD4 as separable inputs required for PMN4-mediated obstruction. Blocking either input reduced gland obstruction without reducing abundance of the PMN4 state, indicating that neutrophil remodeling and obstruction-producing activity are separable. This pattern resembles an AND-gate logic: obstruction proceeds only when both PAD4-dependent histone citrullination and an IFN-γ-dependent requirement are present within the remodeled PMN4 state. Whether IFN-γ and PAD4 act in parallel, in sequence, or through intermediate inflammatory mediators remains unresolved. *Padi4* itself is broadly expressed across neutrophil populations (72), and its transcript was not enriched in PMN4s. Instead, PAD4-dependent activity, like obstruction itself, appears to be governed at the level of execution rather than transcription. Prior work has linked IFN-γ to enhanced NET formation in other inflammatory and tumor contexts (73, 74). Our data suggest a more specific relationship in MGD. Namely, PAD4-dependent histone citrullination can persist during IFN-γ blockade, whereas IFN-γ remains required for obstruction. Thus, IFN-γ is unlikely to act solely by enabling PAD4-dependent citrullination but may instead regulate a distinct step in PMN4-mediated obstruction, either directly or through local secondary mediators. Ex vivo, IFN-γ contributed to acquisition of a PMN4 surface phenotype only alongside a secondary inflammatory signal, consistent with this context-dependent model. Defining the cooperating in vivo signals and the precise relationship between IFN-γ-dependent regulation and PAD4-dependent citrullination will be important next steps.

Together, our findings support immune-mediated obstructive MGD as a mechanistic endotype in which a remodeled neutrophil state forms the cellular basis for IFN-γ- and PAD4-dependent obstruction. The abundance of this remodeled state remains clinically informative and may serve as a biomarker of this MGD endotype, as suggested by our human tear data. However, the murine perturbation studies show that obstruction is determined not by the abundance of this state alone, but by the effector program it is competent to execute. Our work suggests that understanding both the formation of this remodeled neutrophil state and the signals that govern its obstruction-producing activity will be essential for defining and stratifying immune-mediated obstructive MGD.

## METHODS

### Sex as a Biological Variable

For human patient analyses, both male and female patients were included, though equal representation across sexes was not enforced. For murine analyses, published works using the model of interest either recommend or exclusively use female mice (14, 75, 76), and as such most experiments contained here, excluding spatial transcriptomics, also leverage only female mice. Most of our findings have not been evaluated in male mice.

### Patient Recruitment and Human Subjects Approval

This study was conducted in accordance with the tenets of the Declaration of Helsinki and approved by the Duke University Hospital Institutional Review Board (IRB number Pro00068095). Patients presenting to the Duke Foster’s Center for Ocular Immunology were recruited to the study and informed consent was obtained for the prospective collection of ocular surface tear washes as well as chart review. Approved collection of clinical and demographics information including age, sex assigned at birth, and self-identified race was also performed. Inclusion criteria were patients over 18 years of age presenting with non-infectious ocular inflammation without lid abnormalities contributing to ocular surface damage.

### MGD Severity Scoring

A total of 66 consented patients (Table S1) received infrared meibography as part of their normal clinical workup using the Keratograph 5M (Oculus) to evaluate MGD severity before having their tears collected using a sterile saline wash (Gibco, 10010-023) of the ocular surface. Patient MGD severity was qualitatively scored by a clinician based on degree of gland dropout, with severity characterized based on a modified meiboscore (32) scale. Briefly, patients were assigned “Mild” MGD if they exhibited less than 1/3 total gland area reduction, “Moderate” MGD if they exhibited between 1/3 and 2/3 reduction, and “Severe” MGD if they exhibited greater than 2/3 reduction.

### Human Tear Flow Cytometry

Collected tears were stained using a 36-parameter flow cytometry antibody panel and analyzed using a 4-laser Cytek Aurora (UV-V-B-R) as previously described (13, 33) (Table S2). To construct the UMAP and control for batch effect, we included a healthy human blood control sample alongside each tear leukocyte batch. The blood controls were originally collected from a single normal control donor and underwent red blood cell lysis using BD PharmLyse (BD Biosciences, 555899) per manufacturer instructions before being aliquoted and frozen for future use to ensure consistency across sampling days. Sampling days that exhibited batch effect in the control blood samples, characterized by dimensionality reduction results inconsistent with other days, were excluded from downstream analysis. Resulting flow cytometry data was filtered using FlowAI (77) then analyzed using manual gating of leukocytes for cell size, singlets, viability, and CD45 positivity (Fig. S1a) prior to UMAP analysis. UMAP generation included all markers except for linear FSC and SSC markers, markers used in manual gating (CD45, viability dye), or markers which did not obviously stain their established cell populations (CD161, TCRgd, CD20). Downstream leukocyte annotation and filtering of residual debris populations was then performed before heterogeneity analysis. Clustering analysis of all leukocytes was performed using FlowSOM (78) with 30 metaclustering k values and 10 training iterations.

Neutrophils were manually gated (CD16^+^, CD14^-^, CD45RA^-^, CD4^-^, CD3^-^) and the 3 largest clusters (Table S4 – FSOM_K30_01, FSOM_K30_03, FSOM_K30_04) which captured the neutrophil populations were selected for downstream analysis across MGD severities. All flow cytometry analysis of human tear samples was performed using the OMIQ software (Dotmatics).

### Mice

Procedures involving animals were performed with the approval of the Institutional Animal Care and Use Committee at Duke University or AbbVie Inc. and according to approved guidelines. For all experiments excluding spatial transcriptomics, mice were maintained in a specific pathogen free environment at Duke University in cooperation with the Duke University Division of Laboratory Animal Resources. For spatial transcriptomics experiments, mice were maintained at AbbVie Inc. WT C57BL/6 mice (Jax Strain #:000664) were obtained from Jackson Laboratories or bred locally. *Padi4*^-/-^ (Jax Strain #: 030315) and *Ifngr1*^-/-^ (Jax Stain #:003288) mice were bred locally. For the scRNA-seq and spatial transcriptomics experiments only, WT C57BL/6 mice were obtained from Charles River Laboratories (CRL Strain Code 027).

### AED Model

Induction of AED for the spatial transcriptomics experiment is described in its respective section. For all other experiments, MGD was induced in genetically modified and WT female mice aged 6-13 weeks old as previously described (14, 75, 76). Briefly, each mouse received one intraperitoneal (IP) injection containing pertussis toxin (300 ng, Invitrogen, PHZ1174), a solution containing both magnesium hydroxide and aluminum hydroxide (2 mg; Imject Alum, ThermoFisher, 77161 or Regular Strength Antacid Liquid (79), CVS Health, 212532), and ovalbumin (OVA; 10 μg; Sigma, A5503-5G) diluted in PBS (Gibco, 10010-023). Two weeks later, challenges were performed via administration of eyedrops, containing 5 μL of OVA (50 μg/μL) in PBS, to each eye daily for 7 days.

### Murine Meibomian Gland Obstruction Assessment

Meibomian gland orifice obstruction was evaluated as previously described (17, 80). Briefly, mice were lightly anesthetized using ketamine/xylazine (100 μg/gram of body weight, Ketamine Hydrochloride Injection, Dechra, B8U4; 10 μg/gram of body weight, Xylazine, Covetrus, NDC: 11695-4024-1) before having lids gently everted to view the lid margin using a dissecting microscope. Each orifice was qualitatively scored as 1+, 2+, or 3+ based on degree of obstruction, and for each mouse the percentage of orifices scored in each category was calculated to determine MGD severity. For orifice imaging, full thickness eyelids were resected following cardiac perfusion with PBS and placed on ice before imaging the orifices using an Axiocam 208 color camera (Zeiss).

### Tissue Isolation and Processing to Single Cell Suspensions for Flow Cytometry

Whole blood was collected prior to euthanization via cheek puncture of the submandibular vein using a sterile Goldenrod lancet (Medipoint, NC9922361) and placed in an EDTA-coated tube (BD, 365974). Samples were mixed then kept on ice until subsequent processing. Following collection, red blood cell lysis was performed using BD PharmLyse (BD Biosciences, 555899) per manufacturer instructions to generate single cell suspensions for flow cytometry. Tears were collected prior to euthanasia via instillation of 10 μL sterile saline (Gibco, 10010-023) to each eye, followed by immediate recollection. These cells were used as the single cell suspension for flow cytometry. Conjunctiva was collected following CO_2_ euthanasia and transcardiac perfusion. Dissected conjunctival tissue was digested using 1 mg/mL collagenase D (Roche, 11088866001) and 0.05 mg/mL DNase I (Roche, 10104159001) in an HBSS (Gibco 10010-023) solution containing 5% heat inactivated FBS (Gemini Bio-Products, 900-108) and 10 mM HEPES (Gibco, 15630-080) for 45 minutes at 37°C with intermittent vortexing to generate single cell suspensions for flow cytometry. Tarsal plates were collected following CO_2_ euthanasia and transcardiac perfusion with PBS. Dissected eyelids had superficial fascia and conjunctiva resected to isolate the tarsal plate and the contained meibomian glands. Tarsal plates were minced and digested using 1 mg/mL collagenase D (Roche, 11088866001) and 0.05 mg/mL DNase I (Roche, 10104159001) in an HBSS (Gibco 10010-023) solution containing 5% heat inactivated FBS (Gemini Bio-Products, 900-108) and 10 mM HEPES (Gibco, 15630-080) for 45 minutes at 37°C with intermittent vortexing to generate single cell suspensions for flow cytometry.

### Murine Flow Cytometry

Conjunctiva, tarsal plate, and blood single cell suspensions were filtered using 70 μm filters before staining with relevant viability dye and antibody cocktail (Table S9). Stained cells were fixed with stabilizing fixative (BD; 338036) before either being analyzed. The scRNA-seq experiment relied on sorting via the BD FACS Aria III Cell Sorter as described in the scRNA-seq section. Ex vivo neutrophil analysis was performed using a 5-laser Cytek Aurora (UV-V-B-YG-R) with subsequent spectral unmixing and manual gating in FlowJo (BD Biosciences) to identify neutrophils and PMN4s. All other murine samples underwent spectral flow cytometry using a 4-laser Cytek Aurora (UV-V-B-R) with subsequent spectral unmixing, manual gating to isolate neutrophils, and dimensionality reduction analysis in OMIQ (Dotmatics) on the subsetted neutrophils. Neutrophil UMAP generation included the parameters F4/80, CD14, CD62L, PD-L1, CXCR4, ICAM-1, SiglecF, CXCR2, Ly6G, and dcTRAIL-R1. Downstream leukocyte annotation and filtering of residual debris populations was then performed before heterogeneity analysis.

### scRNA-seq Sample Preparation and Cell Sorting

This dataset was generated using 15 9-week-old WT female CRL B6 mice. AED was induced as described above in 9 of these mice, with the remaining 6 mice being left as naive controls that did not receive immunization or challenges. From all mice, blood and conjunctiva were collected and pooled by group, then processed as described above. From AED mice only, tears were isolated as described above and pooled by group. Tear cells were stained with eFluor 450 viability dye (Invitrogen, 65-0863-14) and Calcein AM (ThermoFisher, C1430), blood cells were stained with eFluor 450 viability dye (Invitrogen, 65-0863-14) and CD45 (Biolegend, 103116), and conjunctiva cells were stained with eFluor 450 viability dye (Invitrogen, 65-0863-14), CD45 (Biolegend, 103116), and Ly6G (Biolegend, 127622). Cells were multiplexed by treatment group and by sample type using the Custom BD Single-Cell Multiplexing Set (BD, 626545) per manufacturer instructions. Multiplexed cells were subjected to fluorescence activated cell sorting on a BD FACS Aria III Sorter to enrich for populations of interest by sample type. From blood, only neutrophils were isolated (eFluor 450 viability dye^-^, CD45^+^, Ly6G^+^). From conjunctiva, neutrophils (eFluor 450 viability dye^-^, CD45^+^, Ly6G^+^) and non-neutrophil leukocytes (eFluor 450 viability dye^-^, CD45^+^, Ly6G^-^) were isolated and mixed in a 1:1 ratio. From tears, live cells (Calcein AM^+^, eFluor 450 viability dye^-^) were isolated.

### Library Preparation, Sequencing, and Read Processing

Sorted cells were loaded into the BD Rhapsody and cDNA libraries were prepared using the BD Rhapsody Whole Transcriptome Analysis Amplification Kit per manufacturer instructions. Quality control was performed using the Agilent DNA 4200 Tapestation assay. Libraries were sequenced to a target read depth of ≥50,000 reads per cell by the Duke Sequencing and Genomics Technology Core using the Illumina NextSeq 500 with paired-end sequencing, a 75 base pair read length, 8 bp index, and 20% Phi X. Raw sequencing data was processed using the established Seven Bridges pipeline. Briefly, FASTQ files were generated by demultiplexing and aligned to the mouse genome reference GRCm38-PhiX-gencodevM19-20181206.tar.gz before performing feature barcode processing and unique molecular identifier counting. Resulting counts files were then analyzed using the Seurat package (81) in R.

### scRNA-seq Data Processing and Clustering

ScRNA-seq analysis was performed using the Seurat package in R in accordance with established workflows (81). Briefly, quality control filtering was performed by removing cells with <200 UMI counts or genes that were expressed by less than 3 cells. Cells with <200 or >5500 nFeature, >15% mitochondrial gene percentage, nCount >15000, and multiplexing-indicated doublets were also excluded. After filtering, reads were log normalized and scaled, and the top 2000 variable genes were identified and used to perform principal component analysis (PCA). UMAPs were generated using the top 30 principal components, and clustering was performed using the top 22 principal components. Clusters were annotated using both DEG and canonical cell markers.

### Pseudotime and Pathway Analysis

Pseudotime trajectory analysis was performed on the neutrophil populations using the Monocle3 package (37, 38) as previously described. PMN2 was selected as the root node due to its enrichment in naive mice and in blood, as well as its expression of previously described markers of immature neutrophils (26, 27, 30). fGSEA was performed on the neutrophil populations using the fgsea package (39) with the Mouse Hallmark Gene Set (40, 41) from The Molecular Signatures Database as previously described. DEGs distinguishing PMN4 from PMN1-3 were calculated using the Seurat FindMarkers function, and DEGs were ranked based on adjusted P value and log fold change before performing pathway analysis.

### Cross-Model Gene Module Scoring

Gene module scores were generated as previously described (42) using the top 20-30 DEGs reported to identify neutrophil subpopulations from other murine models of MGD including Pinkie mouse *Lcn2*^lo^ neutrophils (19) and neutrophils enriched in the *Awat2*^-/-^ tarsal plate (22) (Table S8). DEGs that were not present in the AED dataset were excluded.

### Immunofluorescence and Eyelid Imaging

Following CO_2_ euthanasia and transcardiac perfusion with PBS, eyelids were collected and stored on ice. Residual conjunctiva was excised and eyelids were fixed using 4% paraformaldehyde (Santa Cruz Biotechnology, sc-281692) for 1 hour on ice, then serially dehydrated using 15% and 30% sucrose (Sigma-Aldrich, SO389). Dehydrated tissue was embedded in OCT compound (Sakura Finetek, 4583) and frozen at -80°C before being cryosectioned at 12 μm thickness using a Leica CM3050S cryostat. Cryosections were blocked and permeabilized with a solution containing 10% normal donkey serum (Jackson ImmunoResearch, 017-000-121), 0.5% Tween 20 (Sigma, P9416), and 0.5% Triton X-100 (Sigma, T8787) in PBS. Blocked sections were sequentially incubated with primary antibodies and appropriate secondary antibodies with DAPI (Sigma, D8417; 1/1000 dilution) to stain sections. Primary antibodies included Ly6G (Biolegend, 127602; 1/250 dilution), CD14 (ThermoFisher, 60253-1-IG; 1/100 dilution) conjugated with the ThermoFisher Alexa Fluor 594 antibody conjugation kit (ThermoFisher, A88067), and CitH3 (Abcam, ab5103; 1/100 dilution). Secondary antibodies included anti-rat Alexa Fluor 488 (Jackson ImmunoResearch, 712-545-153; 1/500 dilution) and anti-rabbit Alexa Fluor 647 (Jackson ImmunoResearch, 711-605-152; 1/500 dilution). Stained sections were imaged using a Nikon AX R laser scanning confocal system integrated with the Nikon Eclipse Ti2 microscope platform before being analyzed using Fiji (82) or Imaris (Oxford Instruments). Figure panels display maximum intensity projections of stitched large image Z stacks.

### In Vivo Systemic Cytokine Blockade

Systemic antibody-mediated cytokine blockade was performed in AED mice using anti-mouse IFN-γ (BioXCell, BE0055), while associated isotype control mice instead received anti-horseradish peroxidase rat IgG1 (BioXCell, BE0088). Antibodies were diluted to 1 μg/μL in PBS, and 200 μL antibody cocktail was administered to each mouse via intraperitoneal injection, as previously described (52, 53). Intraperitoneal injections began one day before ovalbumin topical challenges and were performed every other day for a total of 4 injections.

### AED Model and Tissue Collection for Spatial Transcriptomics

To generate AED mice, male C57BL/6 mice (9 weeks old; Charles River) received IP immunization on day 0 with 10 μg OVA, 300 ng pertussis toxin, and 4 mg Imject Alum, followed by an IP booster on day 5 with 10 μg OVA and 4 mg Imject Alum. Mice were then challenged topically once daily with 2 μg OVA per eye on days 21–25. Tarsal plate tissues were collected 1 hour after the day 25 challenge from AED and age- and sex-matched control mice. Naïve mice were neither injected nor challenged with OVA.

For tissue collection and processing, eyelids were excised en bloc by gently extending the eyelid with surgical forceps and cutting from corner to corner with surgical scissors. Tissues were oriented conjunctival side down on 3M thick filter paper to minimize curling during fixation in 10% neutral buffered formalin at room temperature for 24 hours, followed by 70% ethanol (catalog number EX0280-3 Supelco, EMD Millipore Corporation) for at least 24 hours (both using 1:20 tissue-to-fixative or alcohol ratio). Samples were then trimmed, placed between biopsy pads in processing cassettes, processed, and then paraffin embedded.

### Sectioning and GeoMx Slide Preparation

For sectioning and GeoMx slide preparation, upper eyelids were serially sectioned horizontally at 5 microns thick and examined microscopically under high-contrast conditions until meibomian glands were present to maximize the number of glands captured within the 35.3 mm × 14.1 mm GeoMx slide capture area. Sections containing the glands were then collected onto GeoMx capture slides. Each slide contained three serial sections. Five consecutive GeoMx slides with the highest or optimal gland content were selected: one for H&E staining, one for GeoMx processing, and three retained as backups. Slides were air dried overnight; all slides except the H&E slide were then immediately stored in slide boxes with desiccant at 4°C until GeoMx processing. The H&E slide was digitally scanned at 20× magnification to assess morphology and select regions of interest in the meibomian glands for GeoMx analysis.

### Tissue Staining and Probe Hybridization

FFPE sections were deparaffinized, rehydrated, and subjected to antigen retrieval and protein digestion as required for assay preparation. Slides were stained with fluorescent morphology markers to delineate tissue compartments. For the whole transcriptome analysis (WTA) panel, markers included panCK (Novus, NBP2-33200, Alexa Fluor 594) and CD45 (Novus, NBP2-34528, Cy5) to distinguish epithelial and immune cell compartments, respectively, with DAPI for nuclear counterstaining. Whole transcriptome RNA probes (Mouse NGS Whole Transcriptome Atlas) were hybridized to the tissue sections.

### ROI Selection and Collection

For ROI selection and collection, slides were scanned on the GeoMx DSP instrument, and regions of interest (ROIs) were selected based on tissue morphology and fluorescent marker signal, targeting anatomical structures including acini and ducts within the meibomian gland. ROIs were further segmented into areas of illumination (AOIs) based on panCK positivity and negativity (e.g., panCK^+^ and panCK^−^ compartments). UV photocleavage was used to release indexing oligonucleotides from each ROI/AOI, which were aspirated and collected into individual wells of a 96-well DSP plate within a 24-hour collection window. Segmented AOIs were collected into separate wells.

### Library Preparation and Sequencing

For library preparation and sequencing, each collected well was converted into an NGS-ready library using the GeoMx SeqCode indexing system (Primer Plate A–H). PCR was performed using 2 μL of collected sample combined with PCR Master Mix and SeqCode primers; all PCR products (4 μL per well) were pooled into a single tube, along with a no-template control (NTC). Pooled libraries were quantified and sequenced on an Illumina NovaSeq 6000 (v1.5 reagents). Sequencing depth was calculated based on total ROI/AOI collection area: WTA required ∼100 reads per μm² of collection area. Sequencing set IDs: A01876:66:HL7W3DRX2.

### Data Processing and Quality Control

For data processing and QC, raw BCL files were converted to FASTQ and demultiplexed using BCLConvert (Illumina) on the internal HPC. FASTQ files were processed through the GeoMx NGS Pipeline v2.3.3.10, encompassing read trimming, stitching, alignment to the mouse genome, and deduplication to generate digital count conversion (DCC) files. Gene-level counts were assembled into an initial dataset containing raw counts with ROI/AOI annotations and sequencing QC metrics.

QC was performed using the GeoMx pipeline. NGS QC flags included NTC counts sum, surface area, nuclei count, deduplicated reads, and negative probe count geometric mean. For WTX data, segment-level QC was applied using a threshold of ≥10% of targets above background; 53 of 58 AOIs passed this filter. Five AOIs were flagged due to nuclei counts below the recommended threshold (∼80–100 nuclei; NanoString cutoff). Sequencing saturation exceeded 50% for all samples, with 92–99% read alignment rates. Negative control probes (Ms IgG1, Ms IgG2a, Rb IgG) and housekeeping positive control targets (GAPDH, Histone H3, S6) were included for normalization evaluation; normalization using negative control background (Rb IgG) was selected based on best correlation as recommended by NanoString.

To perform GeoMx DSP RNA data processing and quality control filtering, raw digital count conversion (DCC) files, the probe configuration (PKC) file, and sample annotations were first imported into R and assembled into a NanoStringGeoMxSet object using the *GeomxTools* and *GeoMxWorkflows* packages. Raw counts were shifted by one to avoid zero values. Segment-level quality control was performed with the following thresholds: a minimum of 1,000 raw reads, ≥80% of reads trimmed, ≥80% of reads stitched, ≥75% of reads aligned, ≥50% sequencing saturation, a minimum negative-control count of 1, a maximum no-template-control (NTC) count of 9,000, a minimum of 20 nuclei, and a minimum segment area of 1,000. Segments failing any criterion were removed. Probe-level quality control (setBioProbeQCFlags) was applied with a minimum probe ratio of 0.1 and a Grubbs outlier failure rate of 20%, and probes flagged as low-ratio or global Grubbs outliers were excluded. Probe counts were then collapsed to gene-level counts (aggregateCounts). The limit of quantification (LOQ) for each segment was defined per probe module as the geometric mean of the negative probes multiplied by their geometric standard deviation raised to the power of 2 (with a floor of 2). Segments with a gene detection rate below 5% were excluded, and genes detected above the LOQ in fewer than 10% of segments were removed. Counts were normalized using the third-quartile (Q3, 75th-percentile) method.

### Clustering and Differential Expression Analysis

The normalized GeoMx object was converted to a Seurat object (as.Seurat, *Seurat*). Variable features were identified, data were scaled, and principal component analysis was performed (30 PCs). A shared-nearest-neighbor graph was constructed on the first 30 principal components, followed by Louvain clustering and UMAP embedding (30 dimensions). Differential expression between groups was assessed with FindMarkers, and p-values were adjusted for multiple testing using the Benjamini–Hochberg (FDR) procedure. Subsequent DEG analysis for each ROI compartment was performed using the heatmaply package (83).

### Gene Set Enrichment Analysis (GSEA)

ssGSEA (84) testing for enrichment of published tarsal plate transcriptomic signatures (22) in our WT AED and naive eyelid spatial transcriptomics dataset was performed using the GSVA R package as previously described. For each group and eyelid region, we used the top 20 published DEGs reported to distinguish the meibocyte and duct cell transcriptomes of healthy B6 mice from those of the *Awat2*^-/-^ MGD model (22) (Table S13). “Healthy” in this analysis indicates use of DEGs reported to exhibit relative upregulation in healthy B6 mice for the respective lid region, while “MGD” indicates use of DEGs reported to exhibit relative upregulation in *Awat2*^-/-^ mice for the respective lid region. GSEA was performed using the Broad Institute’s GSEA software (49, 50) and the Mouse M5 Gene Ontology: Biological Process terms from The Molecular Signatures Database (40) as previously described. For each ROI compartment, DEGs were ranked based on fold change before performing pathway analysis.

### Peritoneal Neutrophil Induction, Isolation, and Ex Vivo Stimulation Assays

Thioglycollate-induced peritonitis and subsequent neutrophil isolation was performed as previously described (57, 58). Briefly, peritonitis was induced via IP injection of 0.5 mL thioglycollate (BD, 221742) in WT C57Bl/6J mice using a 27-gauge needle. Mice were euthanized using CO_2_ 13.5 hours after injection, and peritoneal cells were collected via IP injection of 10 mL sterile DPBS (Gibco, 14190144) using a 27-gauge needle followed by agitation of injected liquid for 1 minute and subsequent recollection of peritoneal exudate using a 19-gauge needle. Neutrophils from individual mice were isolated from peritoneal exudate using a Mouse Neutrophil Isolation Kit (Miltenyi, 130-097-658) per manufacturer instructions.

Isolated neutrophils from individual mice were suspended in an RPMI 1640 media (Gibco, 11875093) solution containing 10% FBS (Gemini, 900-108), 20 mM HEPES buffer (Gibco, 15630080), 2 mM L-glutamine (Gibco, 25030081), and 100 U/mL penicillin-streptomycin (Gibco, 15140122). Approximately 120,000 neutrophils per stimulation condition were isolated from the neutrophil suspension to create paired samples for each biological replicate which were stimulated independently. Neutrophils were stimulated in capped flow cytometry tubes for 3 hours at 37°C using recombinant mouse IFN-γ (100 ng/mL; Gibco, 315-05-100UG), *E. coli* O114 B4 lipopolysaccharide (100 ng/mL; LPS; Sigma-Aldrich, L4391-1MG), or equal volume PBS, as indicated. Stimulated neutrophils were stained for flow cytometry using a subset of antibodies from the PMN4-targeted neutrophil panel (Table S9) sufficient to identify PMN4s and analyzed as described above.

## STATISTICS

Statistical analyses were performed using JMP (SAS) for gland plugging evaluations or R for spatial transcriptomics GSEA, while GraphPad Prism (Dotmatics) was used for all other analyses. Data are presented as means + SEM. Statistical tests used and number of experiment repeats are indicated in the legend of each figure. Shapiro-Wilks normality testing was performed for each analysis and relevant non-parametric statistical tests were used for data that was not normally distributed. All experiments except for sequencing experiments were repeated at least twice.

## STUDY APPROVAL

The human component of this study was approved by the Institutional Review Board at Duke University and written consent was received from patients prior to participation. All animal protocols were approved by the Duke University Institutional Animal Care and Use Committee.

## DATA AVAILABILITY

Murine scRNA-seq and spatial transcriptomics datasets generated by this study will be uploaded to the Gene Expression Omnibus upon publication. Analytic scripts are available upon request.

## AUTHOR CONTRIBUTIONS

Study Design: CJB, DRS, KSH, KRK, VLP ; Data Collection: CJB, SyM, OK, JBC, HMM, RM, DF, JMF, KRK, AA, SL, CY, EMJ, ER, ShM, AAC, AN, JK; Study Management: DRS, VLP, KSH ; Data Analysis: CJB, SyM, OK, JBC, HMM, DF, JMF, KRK, JR, ZW, AA, SL, CY, EMJ, ER, ShM, AAC, AN; Interpretation of Data: CJB, SyM, OK, JBC, DF, JMF, KRK, JR, ZW, AA, SL, CY, EMJ, ShM, AAC, AN, KSH, VLP, DRS ; Manuscript Writing and Narrative Conceptualization: CJB, DRS, OK, SyM ; Review and Approval of Manuscript: CJB, SyM, OK, JBC, HMM, RM, DF, JMF, KRK, JR, ZW, AA, SL, CY, EMJ, ER, ShM, AAC, AN, JK, KSH, VLP, DRS.

## FUNDING SUPPORT

Funding was provided by NIH R01 EY021798 (DRS), NIH P30 EY005722 (DRS/VLP), NIH R01

EY030283 (VLP), NIH R01 EY024484 (VLP), RPB Unrestricted Grant Duke Eye Center (DRS/VLP).

## ACKNOWLEDGMENTS

The authors would like to thank Ashley Moseman, Edward Miao, Mari Shinohara, and Saskia Hemmers for insightful discussions throughout this project. We would also like to thank Edward Miao and Ashley Moseman for providing initial breeder pairs for the *Padi4*^-/-^ and *Ifngr1*^-/-^ mouse lines, respectively. In addition, we would like to thank Teresa Hawks for assistance with patient enrollment. Novel graphical representations were generated using BioRender.com.

## CONFLICT OF INTEREST STATEMENT

C.J.B.: none ; Sy.M.: none ; O.K.: none ; J.B.C.: none ; H.M.M.: none ; R.M.: none ; D.F.: none ; J.M.F.: none ; K.R.K.: AbbVie Inc. - Income, Ownership, Therna Biosciences, LLC. - Income, IP; J.R.: AbbVie Inc. - Income; Z.W.: AbbVie Inc. - Income, Ownership ; A.A.: AbbVie Inc. - Income, Ownership ; S.L.: none ; C.Y.: none ; E.M.J.: none ; E.R.: AbbVie Inc. - Income, Ownership ; Sh.M.: AbbVie Inc. - Income, Ownership ; A.A.C.: AbbVie Inc. - Income, Ownership ; A.N. - AbbVie Inc. - Income, Ownership ; J.K.: none ; K.S.H.: AbbVie Inc. - Income, Ownership ; V.L.P.: B&L: consultant, BrightStar: consultant, Brill Engine: consultant, BRIM: advisory board, Biocryst: consultant, Dompe: consultant, EmmeCell: advisory board, NIH/NEI: grant support, Oculis: consultant, Ocubio: Co-Founder, ORA: consultant, Senjun: consultant, Tarsus: consultant, Tearsolutions: research support, Thea: consultant, Trefoil: equity and advisory board ; D.R.S.: none.

---

**Figure S1.**
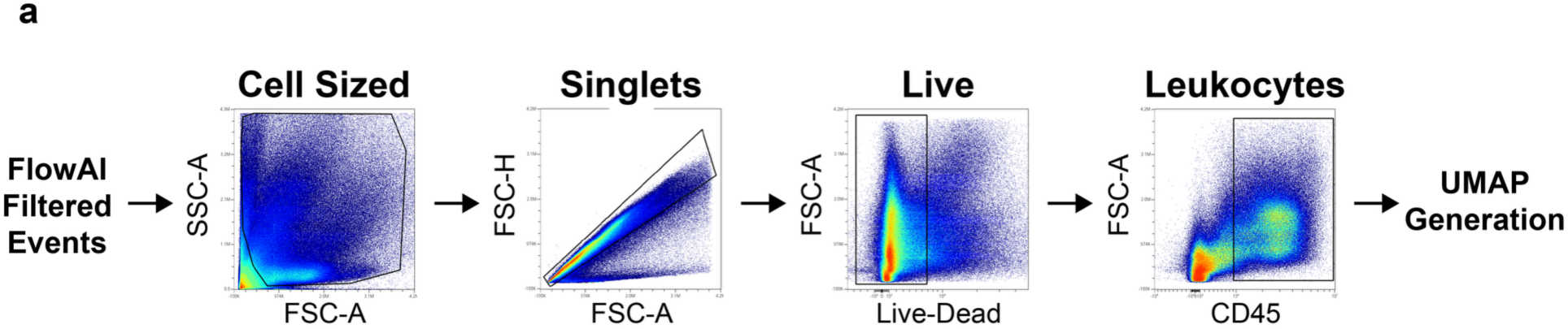
Evaluating human tear leukocytes using 36-parameter spectral flow cytometry. **(A)** Manual flow cytometry gating strategy used to isolate human CD45^+^ leukocytes for downstream dimensionality reduction analysis.

**Figure S2.**
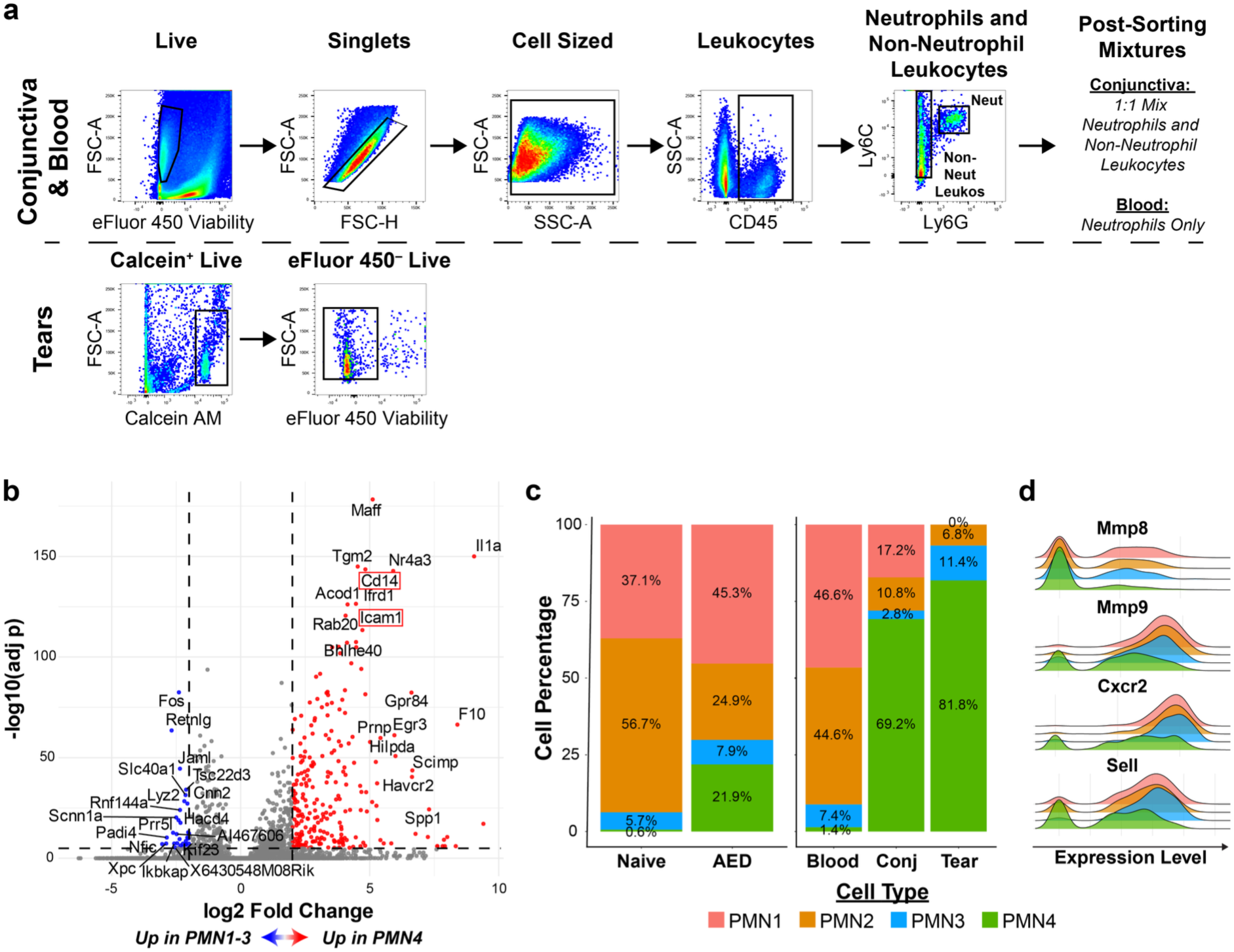
ScRNA-seq gating strategy and neutrophil analysis supporting data. **(A)** FACS gating strategy used in scRNA-seq to enrich for specific cell populations from conjunctiva, tears, and blood from WT AED and naive mice. **(B)** Volcano plot showing DEGs which distinguish PMN1-3 from PMN4. Red boxes around *Cd14* and *Icam1* highlight key genes used to identify PMN4s downstream in protein analyses. **(C)** scRNA-seq quantification of the percentage of neutrophils clusters in each disease state and type of sample, respectively. **(D)** RidgePlots displaying expression of key blood- and neutrophil immaturity-associated markers for each neutrophil cluster.

**Figure S3.**
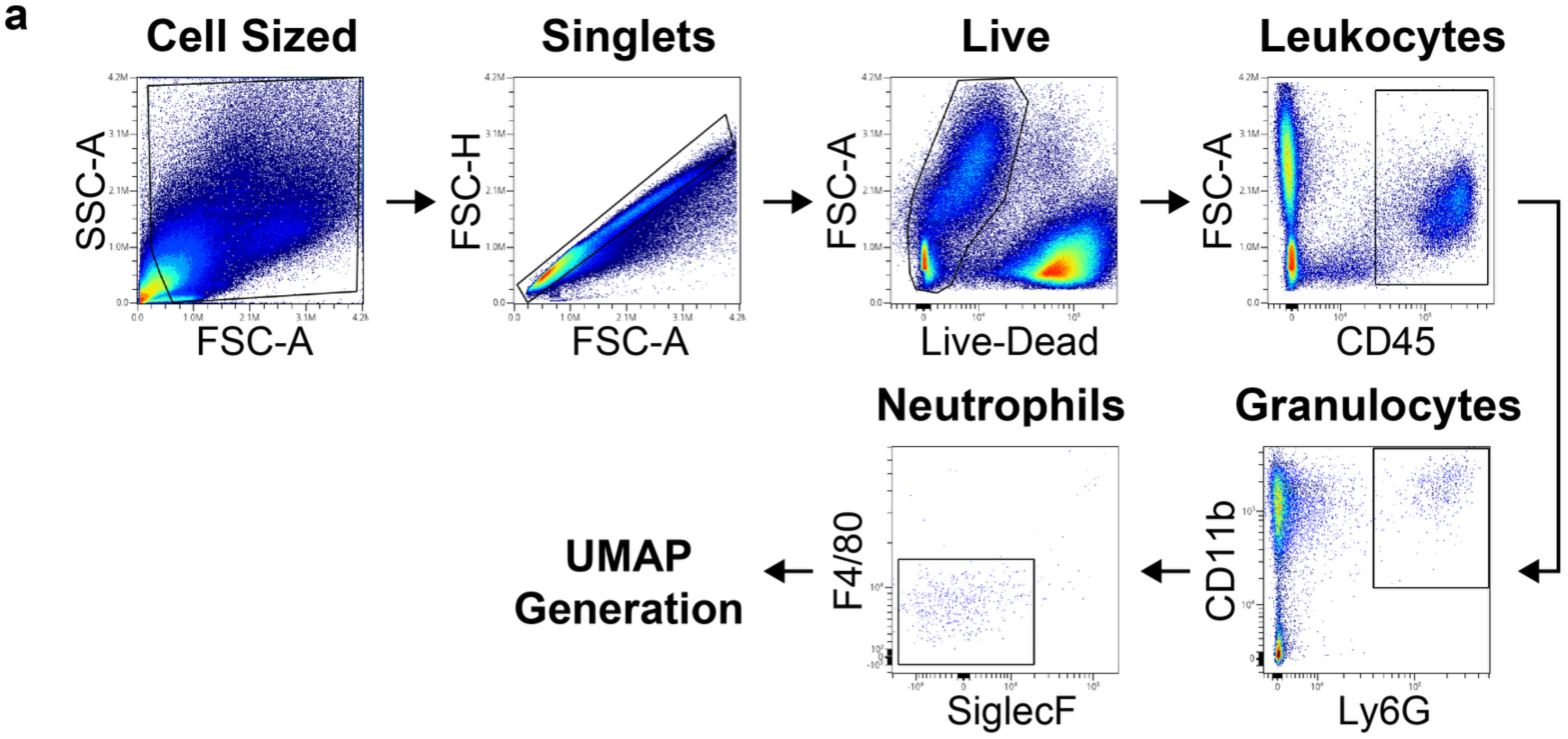
Strategy for analyzing spectral flow cytometry in murine samples. **(A)** Gating strategy used in murine flow cytometry experiments for dimensionality reduction analyses.

**Figure S4.**
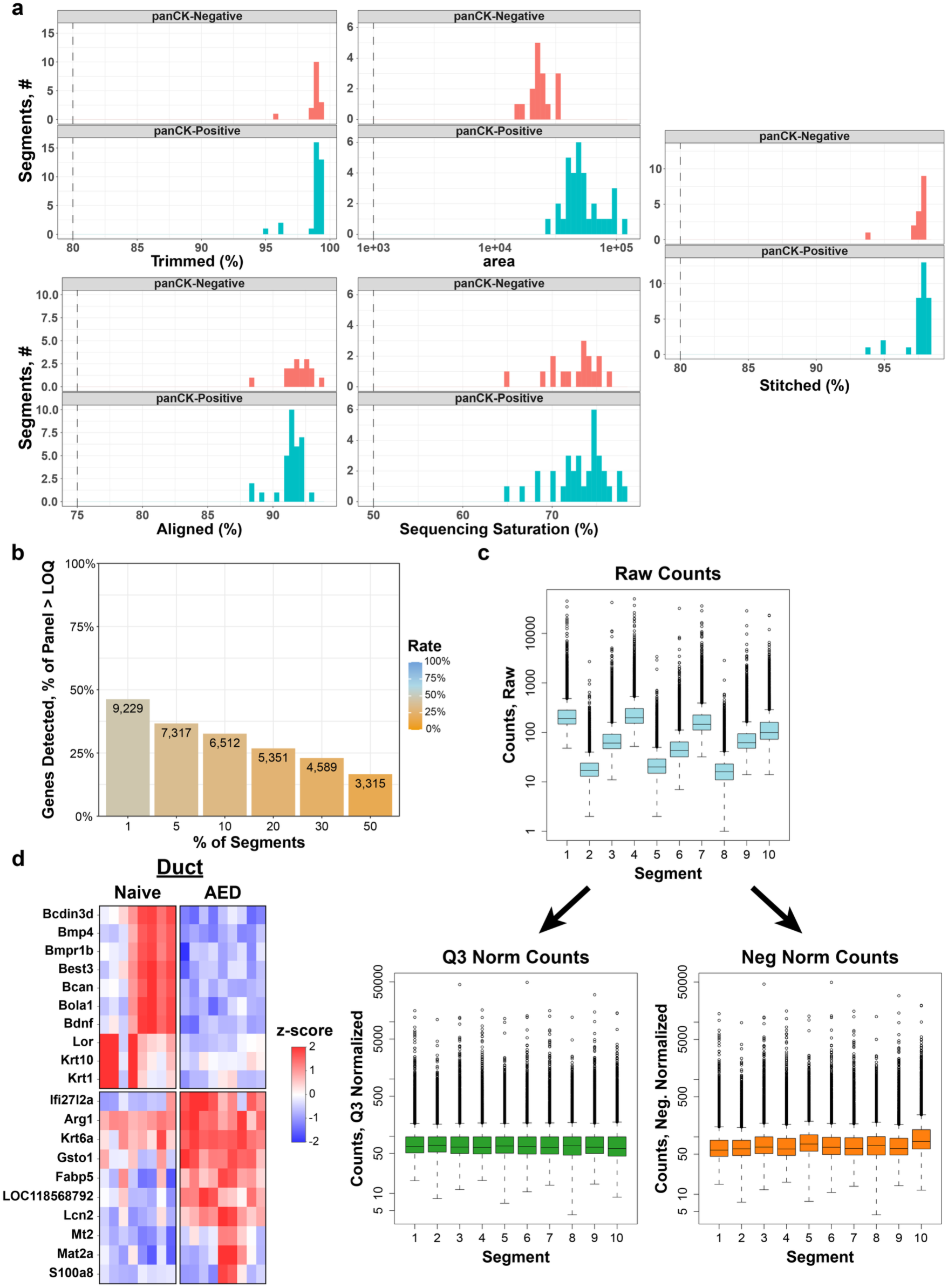
Spatial transcriptomics QC metrics and duct DEGs between WT naive and AED mice. **(A)** Key QC metrics for panCK-positive and panCK-negative segments demonstrating high dataset quality. **(B)** QC-focused quantification of genes detected across segments demonstrating high dataset quality. **(C)** Evaluating Q3 and negative control normalization strategies supporting use of Q3 normalization. **(D)** Heatmap showing DEGs distinguishing WT naive and AED duct ROIs.

**Figure S5.**
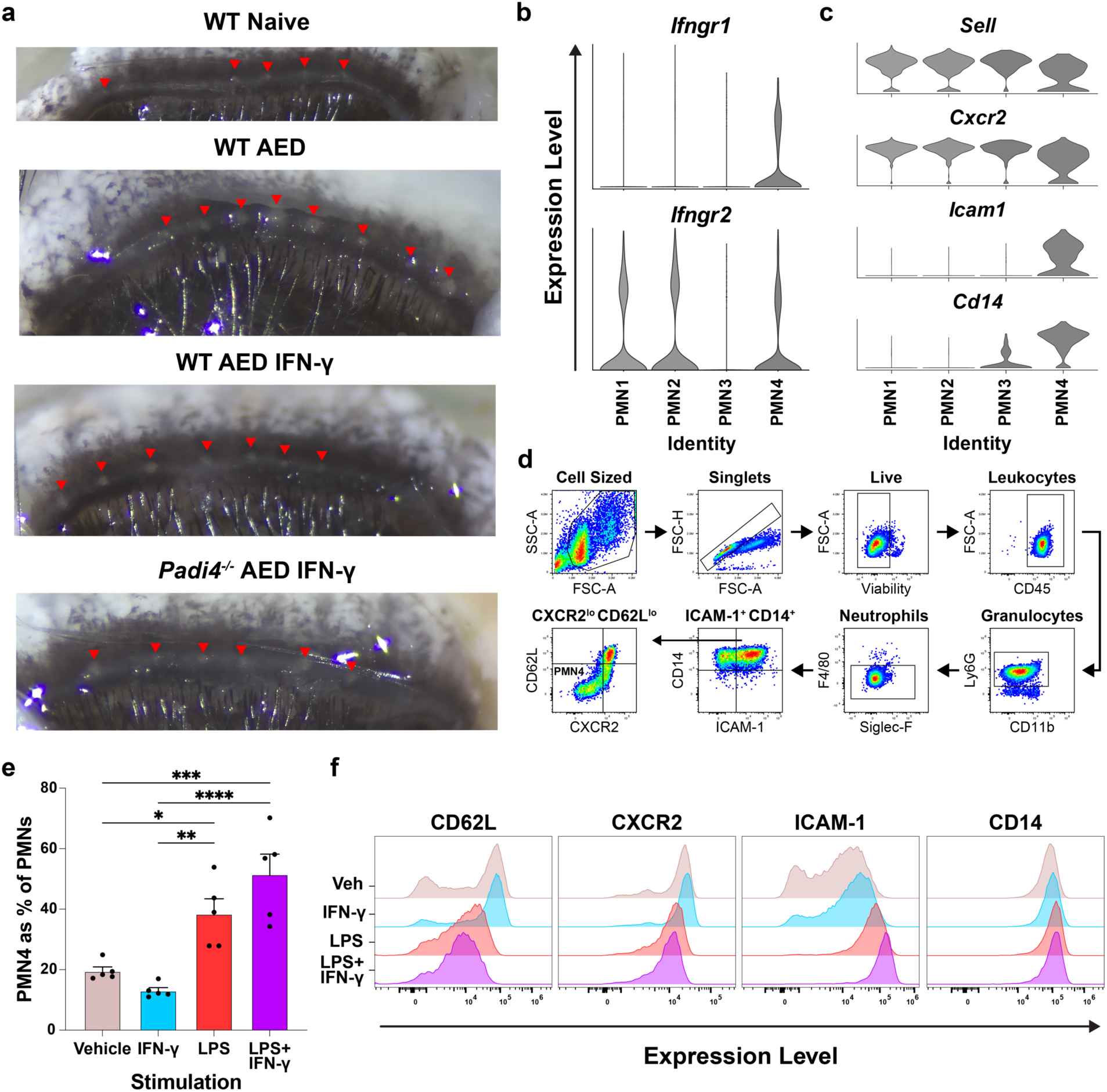
IFN-γ is necessary for MG orifice obstruction and cooperates with inflammatory factors to induce a PMN4 phenotype. **(A)** Representative gland orifice images showing reduction in AED-induced orifice obstruction with IFN-γ blockade and *Padi4^-/-^.* Red arrows highlight gland orifices. **(B)** scRNA-seq data showing expression of IFN-γ receptor heterodimer subunits in PMN1-4. **(C)** scRNA-seq data showing expression of key PMN4 markers used to identify PMN4s in the reductionist ex vivo stimulation assay. **(D)** Manual gating strategy used to identify PMN4s in the thioglycollate-induced peritoneal neutrophil ex vivo stimulation experiments. **(E)** Quantification of flow cytometry data from thioglycollate-induced peritoneal neutrophils stimulated ex vivo for 3 hours (N=5 biological replicates per condition). Dots represent quantification of each biological replicate. Values represent the percentage of all neutrophils that exhibited a PMN4 phenotype (CD14^+^ ICAM-1^+^ CD62L^lo^ CXCR2^lo^) after stimulation. **(F)** Representative flow cytometry histograms of key PMN4-associated marker expression (CD14, ICAM-1, CD62L, and CXCR2) in ex vivo stimulated, thioglycollate-induced peritoneal neutrophils. Data were collected from 2-3 independent experiments. *: P < 0.05 ; **: P < 0.01; ****: P <0 .0001 ; ns: not significant. ANOVA with Tukey’s multiple comparison post-hoc test (E). Plots show mean + SEM.

